# Cognitive Aging in the Rhesus Monkey: A Comprehensive Multi-Domain Analysis Across the Adult Lifespan

**DOI:** 10.64898/2026.11.18.767608

**Authors:** Anna W. Ganzabal, Tara L. Moore, Jon Rueckemann, Ana Vitantonio, Ronald J. Killiany, Mark B. Moss, Sara Ibañez, Klaus Wimmer, Douglas L. Rosene

**Author notes:** Corresponding Author: Tara L. Moore, Ph.D., Boston University Chobanian & Avedisian School of Medicine 700 Albany Street, W701, Boston, MA 02118. Ganzabal, AW and Moore, TL – co-1st authors Ibañez, S, Klaus Wimmer^C^, and Rosene DL – co-senior authors.

## Abstract

In contrast to neurodegenerative aging as in Alzheimer’s Dementia (AD), normal cognitive aging in humans is characterized by gradual, progressive declines in learning, memory, and executive function. Yet the precise timing and trajectory of these changes remain difficult to establish due to confounds including subclinical pathology from early AD or other neurodegenerative diseases. The rhesus monkey, which shares close neurobiological homology with humans but does not develop dementia, offers a powerful model for studying normal cognitive aging. Here, we present a comprehensive analysis of cognitive performance across the adult lifespan in 99 male and female rhesus monkeys (ages 4.5–32 years), including a critically underrepresented middle-aged group, tested on tasks of recognition memory, spatial and object working memory and executive function. Continuous-age regression analyses revealed graded declines across multiple aspects of cognitive performance. Increasing age was associated with poorer rule acquisition, reduced working-memory capacity, diminished practice-related improvement, and impaired abstraction and set-shifting, although the strength and timing of these relationships varied across measures. Categorical comparisons provided a coarser but complementary view of these patterns with differences most often distinguishing young from aged animals, although selected measures, also differentiated middle-aged animals, specifically, improvement on the spatial working-memory task and perseverative responding. Learning-curve analyses showed that the ability to improve with experience declined with age. No significant sex differences were observed.

**Highlights:**

Cognitive aging in monkeys is expressed as a gradual shift in population-level performance
Age predicts reduced cognitive capacity and a diminished ability to improve performance
Cognitive aging is not expressed uniformly across domains
Progressive impairments in rule learning, working memory, and cognitive flexibility with age
Cognitive aging in monkeys closely mirrors normal human aging across multiple domains

## Introduction

Normal cognitive aging is characterized by progressive but highly variable changes in learning, memory, and executive function. In humans, the timing and severity of these changes differ substantially across individuals, and the earliest stages of decline can be difficult to distinguish from mild cognitive impairment, incipient dementia, or the cumulative effects of health, lifestyle, and environmental exposures. These confounds make it challenging to isolate the pattern of normal cognitive aging and to determine whether age-related changes emerge abruptly at a particular stage of adulthood or develop progressively across the lifespan.

The rhesus monkey provides a powerful model for addressing this question. Rhesus monkeys share important neuroanatomical and cognitive similarities with humans but do not develop the dementing illnesses that commonly complicate studies of human aging. Prior studies have established age-related impairments across multiple cognitive domains, including recognition memory, working memory, and executive function.^1–18^ Although large-cohort studies have convincingly demonstrated cognitive differences between young and aged rhesus monkeys across multiple domains, ^4^ the trajectory of change across the intervening years remains less well characterized.

Middle age is therefore an important but underrepresented interval in studies of cognitive aging. In humans, measurable cognitive changes begin during middle age, alongside alterations in synaptic function, myelin integrity, and inflammatory signaling.^19^ Studies in rhesus monkeys similarly suggest that impairments in executive function and spatial working memory can emerge during middle age.^8^ However, to date studies of middle-aged cohorts have been relatively small and have limited the ability to determine whether these changes reflect a distinct transition or form part of a broader continuum of age-related decline.

The present study addresses this gap by characterizing cognitive performance in a large cohort of male and female rhesus monkeys sampled across the adult lifespan. We examined complementary cognitive capacities using three behavioral tasks: recognition memory and rule acquisition using the Delayed Non-Matching to Sample task; spatial and object working-memory capacity using the Delayed Recognition Span Tasks; and abstraction and set-shifting using the Category Set Shifting Task. By analyzing age both as a continuous variable and augmenting with conventional age-group comparisons, we sought to characterize progressive lifespan relationships that were previously only detectable between broad age categories. This approach provides a more detailed account of the trajectory of cognitive aging in the rhesus monkey and establishes a normative foundation for identifying the neurobiological mechanisms that contribute to early cognitive decline.

## Methods

### Subjects

Cognitive data for this study was collected from 99 rhesus monkeys (Macaca mulatta) of both sexes between the ages of 4.5 and 32 years old using identical testing protocols. Out of these 99 monkeys, 82 monkeys completed all tasks (Table 1). Based on an extensive survival study at Emory Regional Primate Research Center, which suggests a ratio of three to one between monkey and human years of age, we have classified young monkeys as between 5 and 13 years of age, middle-aged monkeys as between 13 and 20 years of age and aged monkeys as > 20 years old.^20^ As detailed in Tables 1 and 2, our dataset is well balanced in terms of tasks, age groups and sex. Note that one monkey aged into the middle-aged group during testing and six monkeys aged into the aged group. All monkeys had known birth dates, complete health records and were obtained from National Primate Research Centers or private vendors. All monkeys received medical examinations before entering the study. In addition, explicit criteria were used to exclude monkeys with a history of any of the following: splenectomy, thymectomy, exposure to radiation, cancer, organ transplantation, malnutrition, chronic illness including viral or parasitic infections, neurological diseases or chronic drug administration. Each of the monkeys underwent magnetic resonance imaging (MRI) to ensure there was no occult neurological damage. Results of the evaluations revealed that all monkeys were healthy when they were entered into the study.

**Table 1.** List of monkeys sex, age group, age at testing and conditions included in analysis. Age groups: Y – young, MA – middle-aged, A – aged.

| ID | Sex | Age Group | Age at DNMS | Age at DRST | Age at CSST |
|---|---|---|---|---|---|
| **AM095** | F | Y | 6.0 | 6.2 | 7.0 |
| **AM222** | M | Y | 7.4 | 6.3 | 72 |
| **AM204** | M | Y | 4.5 | 6.3 | 5.9 |
| **AM358** | F | Y | 6.2 | 6.4 | n/a |
| **AM230HL** | M | Y | 6.4 | 6.6 | 6.8 |
| **AM280** | M | Y | 6.5 | 6.9 | 7.8 |
| **AM229HL** | M | Y | 6.8 | 6.9 | 7.1 |
| **AM132** | M | Y | n/a | n/a | 7.2 |
| **AM290** | F | Y | 6.8 | 7.1 | n/a |
| **AM368** | M | Y | 6.9 | 7.2 | 7.6 |
| **AM255** | F | Y | 7.5 | 7.3 | 10.0 |
| **AM299** | F | Y | 7.1 | 7.3 | n/a |
| **AM367** | M | Y | 7.1 | 7.6 | 8.2 |
| **AM296** | M | Y | 7.7 | 7.9 | 8.4 |
| **AM289** | M | Y | 7.9 | 8.2 | 8.6 |
| **AM254** | F | Y | 8.9 | 9.1 | 10.7 |
| **AM163** | F | Y | 8.7 | 9.1 | 9.7 |
| **AM352** | M | Y | 8.9 | 9.2 | 9.4 |
| **AM214** | F | Y | 8.8 | 9.3 | 10.1 |
| **AM202** | F | Y | 9.1 | 9.4 | 10.1 |
| **AM197** | F | Y | 9.2 | 9.8 | n/a |
| **AM295** | M | Y | 9.5 | 9.9 | 10.4 |
| **AM195** | F | Y | 10.5 | 10.9 | 11.8 |
| **AM303** | M | Y / MA | 11.9 | 12.3 | 12.5 |
| **AM291** | F | MA | 12.7 | 13.0 | 13.3 |
| **AM278** | F | MA | 13.2 | 13.7 | 14.8 |
| **AM353** | M | MA | 13.9 | 14.0 | 14.3 |
| **AM034L** | M | MA | n/a | n/a | 15.2 |
| **AM143** | M | MA | 14.5 | 14.7 | 15.5 |
| **AM288** | M | MA | 14.6 | 15.1 | 15.6 |
| **AM340** | F | MA | 14.8 | 15.1 | 15.2 |
| **AM257** | F | MA | 15.0 | 15.2 | 17.0 |
| **AM285** | F | MA | 14.4 | 15.3 | 16.4 |
| **AM158h** | M | MA | 15.5 | 15.7 | 16.9 |
| **AM253** | F | MA | 16.0 | 16.4 | 17.9 |
| **AM251** | F | MA | 16.1 | 16.5 | 16.7 |
| **AM274** | M | MA | 16.2 | 16.6 | 17.7 |
| **AM250** | F | MA | 16.1 | 16.6 | 16.9 |
| **AM223L** | M | MA | 16.5 | 16.8 | 17.5 |
| **AM297** | M | MA | 16.6 | 17.0 | 17.4 |
| **AM233H** | M | MA | n/a | n/a | 17.9 |
| **AM252** | F | MA | 17.0 | 17.2 | 18.8 |
| **AM190** | F | MA | 16.6 | 17.2 | 17.8 |
| **AM300** | F | MA | 17.4 | 17.6 | 18.2 |
| **AM209L** | M | MA | 16.9 | 18.1 | 18.2 |
| **AM161** | F | MA | 17.8 | 18.2 | 18.7 |
| **AM149** | F | MA | n/a | n/a | 19,2 |
| **AM256** | F | MA | 18.0 | 18.4 | 19.9 |
| **AM159** | F | MA | 18.2 | 18.6 | 19.4 |
| **AM293** | F | MA | 18.3 | 18.7 | 19.2 |
| **AM124** | M | MA | 18.7 | 19.0 | 19.2 |
| **AM133** | M | MA | 19.1 | 19.0 | 19.2 |
| **AM226L** | M | MA | 18.8 | 19.0 | 19.9 |
| **AM212** | F | MA | 18.9 | 19.3 | 20.0 |
| **AM390** | M | MA | 19.1 | 19.4 | 19.7 |
| **AM391** | M | MA | 19.1 | 19.4 | 19.7 |
| **AM160** | F | MA / A | 18.2 | 19.6 | 20.5 |
| **AM372** | M | MA / A | 19.2 | 19.7 | 20.3 |
| **AM314x** | M | MA / A | 19.7 | 20.0 | 20.2 |
| **AM123** | M | MA / A | n/a | n/a | 20.4 |
| **AM126** | F | MA / A | 19.8 | 20.1 | 20.4 |
| **AM287** | F | MA / A | 19.8 | 20.2 | 20.6 |
| **AM286** | F | A | 20.2 | 20.7 | 21.5 |
| **AM242** | M | A | 20.5 | 20.8 | 20.5 |
| **AM236** | M | A | 20.5 | 20.9 | 21.5 |
| **AM162** | F | A | 20.6 | 21.0 | 22.0 |
| **AM196** | F | A | 20.8 | 21.2 | 22.3 |
| **AM281** | M | A | n/a | n/a | 22.5 |
| **AM208** | M | A | 21.1 | 21.4 | 22.5 |
| **AM234** | F | A | 21.7 | 22.2 | 23.0 |
| **AM282** | M | A | 21.8 | 22.4 | 22.6 |
| **AM389** | F | A | 22.1 | 22.6 | 22.9 |
| **AM395** | F | A | 22.2 | 22.6 | 22.9 |
| **AM276** | M | A | 22.4 | 22.7 | 23.6 |
| **AM179** | F | A | 22.2 | 22.8 | 23.2 |
| **AM356** | M | A | 22.9 | 23.3 | 23.7 |
| **AM243** | M | A | n/a | n/a | 24.0 |
| **AM090** | F | A | 23.5 | 23.6 | 24.4 |
| **AM235** | F | A | 22.9 | 23.6 | 24.2 |
| **AM284** | M | A | 23.8 | 24.3 | 25.1 |
| **AM283** | M | A | 24.8 | 25.3 | 26.1 |
| **AM374** | M | A | 24.9 | 25.3 | 26.0 |
| **AM369** | F | A | 25.1 | 25.4 | n/a |
| **AM301** | M | A | 25.2 | 25.4 | n/a |
| **AM371** | F | A | 25.2 | 25.5 | 25.8 |
| **AM406** | F | A | 25.2 | 25.5 | 25.7 |
| **AM109** | M | A | n/a | n/a | 26.4 |
| **AM 298** | M | A | 25.5 | 25.9 | 26.5 |
| **AM304** | M | A | 25.5 | 25.9 | 26.1 |
| **AM275** | F | A | 25.9 | 26.3 | 27.1 |
| **AM370** | F | A | 26.2 | 26.6 | 27.2 |
| **AM355** | M | A | 26.5 | 26.9 | 27.1 |
| **AM098** | F | A | 26.6 | 26.9 | 27.7 |
| **AM210** | F | A | 27.1 | 27.7 | n/a |
| **AM180** | F | A | 28.5 | 28.8 | n/a |
| **AM024L** | F | A | n/a | n/a | 29.4 |
| **AM373** | M | A | 28.9 | 29.6 | 29.8 |
| **AM091** | M | A | 28.6 | 29.7 | 30.0 |
| **AM402** | M | A | 32.0 | 32.2 | 32.4 |

**Table 2.** Total number of monkeys included in each task. divided by sex and age group.

|  |  | **DNMS** | **DRST** | **CSST** |
|---|---|---|---|---|
| Young | Female | 11 | 11 | 7 |
|  | Male | 12 | 11 | 12 |
| Middle-aged | Female | 19 | 17 | 17 |
|  | Male | 15 | 15 | 15 |
| Aged | Female | 17 | 19 | 19 |
|  | Male | 16 | 17 | 21 |
| **Total Count** |  | 90 | 90 | 91 |

While on study, monkeys were individually housed in colony rooms in the Boston University Animal Science Center where they were in constant auditory and visual range of other monkeys. This facility is fully AAALAC accredited, and animal maintenance and research were conducted in accordance with the guidelines of the National Institutes of Health and the Institute of Laboratory Animal Resources Guide for the Care and Use of Laboratory Animals. All procedures were approved by the Boston University Institutional Animal Care and Use Committee. Diet consisted of Lab Diet Monkey Chow (#5038 - LabDiet Inc., St. Louis, MO) supplemented by fruit and vegetables with feeding taking place once per day, immediately following behavioral testing. All monkeys were fed 12–20 biscuits per day per day to maintain a stable body weight. During testing, small pieces of fruit or candy were used as rewards. Water was available continuously. The monkeys were housed under a 12-h light/dark cycle with cycle changes occurring in a graded fashion over the course of an hour. They were checked daily by trained observers for health and well-being and were given a medical exam every three months by a Clinical Veterinarian in the Boston University Animal Science Center.

### Cognitive Testing

The behavioral battery assessed recognition memory, spatial and object working memory capacity, and executive functions related to abstraction and set shifting. In addition, we also quantified rule acquisition and the capacity to improve with practice. As detailed in Table 1, not all monkeys completed every task. Because some monkeys completed different tasks at different ages, their age-group classification could vary across tasks (Table 2). For categorical analyses, each monkey was assigned to the age group corresponding to their age when the relevant task was performed.

Monkeys were initially familiarized with behavioral testing in a Wisconsin General Testing Apparatus (WGTA) and trained to displace a single gray plaque placed pseudo-randomly over one of the three food wells to obtain a reward. Monkeys were trained until they responded for 40 consecutive trials on one day. Then monkeys completed the Delayed Non-Matching to Sample (DNMS) task, the Delayed Recognition Span Tasks (DRST) and the Category Set Shifting Task (CSST). For all tasks, For all tasks, the testing apparatus had a darkened interior and was in a darkened room and white noise was played on two speakers located within the automated apparatus to mask extraneous sounds. A non-correctional procedure was used, such that an incorrect response was recorded without repeating the trial. Small pieces of candy or fruit were used as rewards.

### Delayed Non-Match to Sample (DNMS)

The DNMS task assesses recognition memory by requiring monkeys to distinguish a novel object from a recently presented sample object. The first phase of testing assesses learning the non-match rule until criterion performance was reached (90% correct over 100 trials). Asymptotic recognition memory performance was then assessed by giving a fixed number of trials with increased delays between sample presentation and the choice first to 120 and then to 600 seconds.

### DNMS Acquisition

As shown in **Figure 1A**, each trial begins with a sample object presented over the central baited food well. The monkey was permitted to displace the object and obtain the reward. The door was then lowered, and the original, now familiar, sample object was placed over an unbaited lateral well and a new, novel, unfamiliar object is placed over the other lateral well that was baited. Ten seconds after the original sample trial, the choice phase began, and the monkey must choose the unfamiliar, novel object to obtain the reward. Twenty seconds later, the next trial was initiated with novel objects. The position of the two objects on successive recognition trials was varied from left to right lateral wells in a pre-determined pseudorandom order. A non- correction procedure was used and thirty trials per day were given until the monkey reached a learning criterion of 90 correct responses in 100 consecutive trials or a maximum of 1500 trials. Objects were drawn from a pool of 600 "toy" objects and paired so that in each daily session of 30 trials, 60 of the objects were used. Once all of the initial pairings were used, (25 days of 30 trials per day), the 600 objects were randomly recombined to produce new pairs so that the pairings presented continued to be new and unique on each trial.

**Figure 1.**
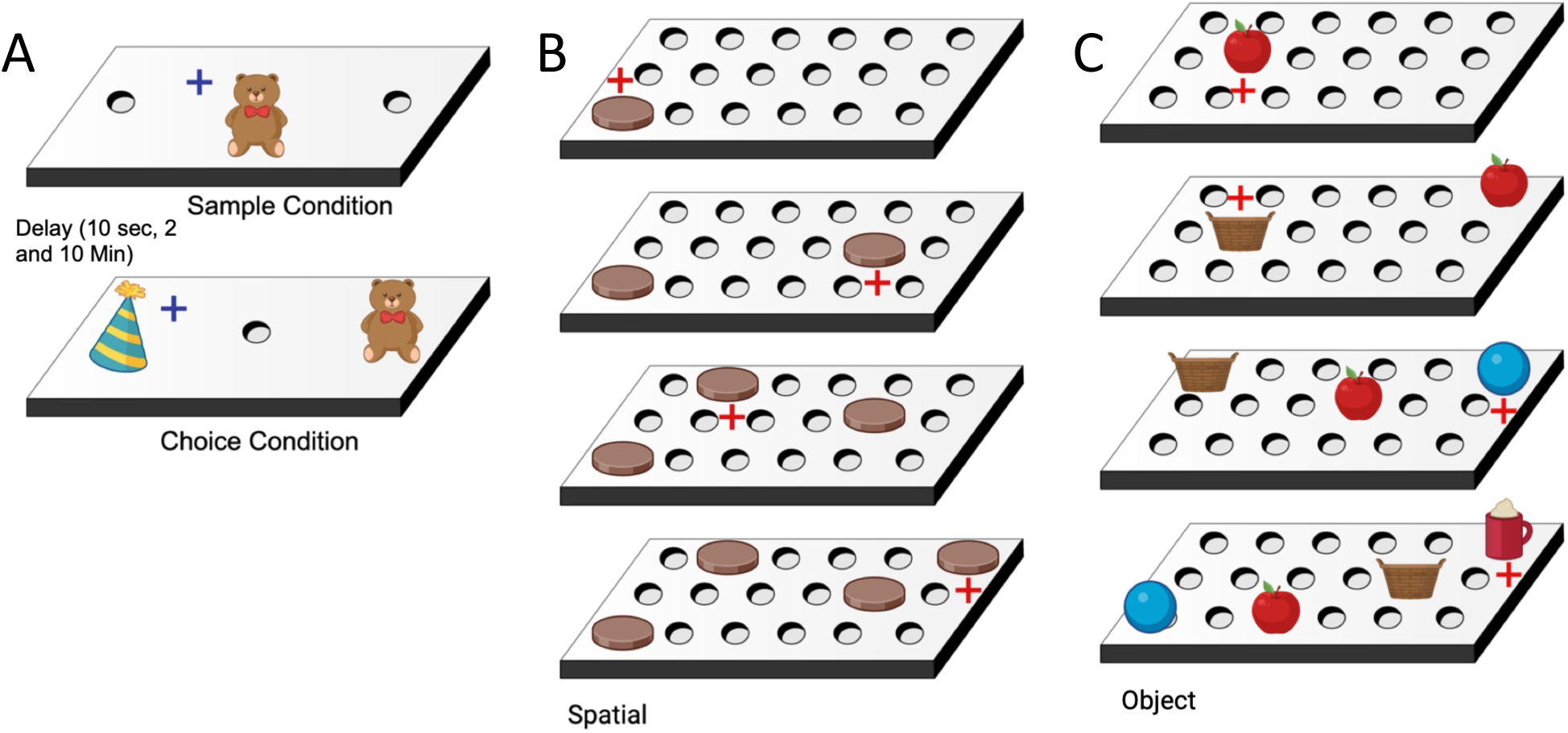
A) A schematic of the Delayed Non-Match to Sample showing a sample object presented over the central baited food well. The monkey is permitted to displace the object and obtain the reward. The door is then lowered, and the original, now familiar, sample object is placed over an unbaited lateral well and a novel, unfamiliar object is placed over the other lateral well that is baited. Ten seconds (or 2 or 10 minutes during the delay condition) after the original sample trial, the choice trial begins and the monkey is required to choose the unfamiliar, novel object in order to obtain the reward. Following a 10-s inter-trial interval, the next trial is initiated with two new objects. B) Schematic of the Delayed Recognition Span Task - Spatial showing sequential stimuli (brown disc) presentation within one trial of the DRST. During each presentation the monkey must choose the disc in the new spatial location. Each successive correct response trial was followed by the addition of a new disc in a novel location on the testing board, and this continued until the monkey made an error (i.e. chose a previously chosen disc). With the occurrence of the first error, the trial was terminated, and the number of discs on the testing board minus one was counted to determine the recognition span score for that trial (i.e. number of correct consecutive responses). C) Schematic of the Delayed Recognition Span Task – Object showing sequential stimuli (small toy) presentation within one trial of the DRST. During each presentation the monkey must choose the new toy. Each successive correct response trial was followed by the addition of a new toy in a novel location on the testing board, and this continued until the monkey made an error (i.e. chose a previously chosen disc). With the occurrence of the first error, the trial was terminated, and the number of discs on the testing board minus one was counted to determine the recognition span score for that trial (i.e. number of correct consecutive responses).

### DNMS Delays

Following acquisition of the basic task, the 10 sec delay between the presentation of the sample object and the recognition trial was increased, in stages, first to 120 seconds and then to 600 seconds. Ten trials a day for 10 days at each delay interval were given with the monkey remaining in the testing apparatus during the delay interval. A total of 100 trials were given over ten days at each of the two delays.

### Delayed Recognition Span Task-Spatial (DRST-sp)

The spatial version of the Delayed Recognition Span Task (DRSTsp) assesses spatial working memory capacity under pressure of accumulating interference. As shown in **Figure 1B**, the DRST testing board has three rows of six wells each (3.5 cm wide, 0.5 cm deep) with the wells spaced 6 cm apart within a row and the rows spaced 1.5 cm apart. For this task, the stimuli were 15 identical plain brown discs (6 cm in diameter). A trial consisted of multiple stimulus sequences presented sequentially until an error was made. During the first sequence of a trial, with the testing board obscured by an opaque panel, one disc was placed over 1 of the 18 wells and baited with a food reward. The WGTA door was raised, and the monkey was allowed to displace the disc to obtain the reward. The door was then lowered, the first disc was returned to its original position over the now unbaited well, and a second disc was placed on the board over a baited well in a different spatial location. After 10 s, the door was once again raised, and the monkey was required to identify the new second disc in its novel spatial location to obtain the reward. Each successive correct response trial was followed by the addition of a new disc in a novel spatial location on the testing board over a newly baited well and this continued until the monkey made an error (i.e., chose a previously chosen disc). With the occurrence of the first error, the trial was terminated and the number of discs on the testing board minus one were counted to determine the recognition span score for that trial (i.e., number of correct consecutive responses). Ten such trials were presented each day, 7 or 8 of which used unique spatial location of discs and 2 or 3 of which used a repeated series of spatial locations of discs, for 10 days (total 100 trials).

### Delayed Recognition Span Task-Object (DRST-ob)

The object version of Delayed Recognition Span Task (DRSTob) assessed object working memory capacity under increasing interference independent of spatial cues. As shown in **Figure 1C**, the testing board for DRSTob was the same as for DRSTsp. The object condition of the DRST was administered in the same manner as the spatial condition except the stimuli consisted of trial unique toy objects. As with DRSTsp, the trial began with a single object covering a baited well. Once that object was displaced and reward retrieved the previously presented object was placed over a new unbaited well and new object was placed over a newly baited well. When the door was raised the subject had to identify the new object regardless of location. Each new object was placed over a new location, and all of the previous objects were moved to new locations to preclude the use of spatial cues. This continued until an error was made. Ten such trials were presented each day, 7 or 8 of which used a unique set of objects and 2 or 3 of which used a repeated set of objects, for 10 days (total 100 trials).

### Category Set Shifting Task (CSST)

The CSST assessed abstraction and set shifting by requiring monkeys to learn and flexibly update rewarded stimulus categories based on accumulated evidence. Testing progressed in three stages: an automated pretraining to accustom the monkey to responding to an object on the touch screen, a three-choice discrimination task followed by formal CSST testing. Both stages were administered in the same but now automated WGTA containing a touch-sensitive monitor driven by a Macintosh computer (1.83 GHz Intel Core 2 Duo Processor). An automated pre-training task was used to teach the monkey to touch the computer screen. The pre-training task required the monkey to touch a single stimulus which appeared in one of 9 random locations on the screen to receive a food reward. This procedure was continued for 20 trials a day until the monkey correctly responded for all 20 trials. If a monkey did not respond within one minute, the screen reverted to black, a non-response was recorded, and the intertrial interval began followed by the next trial. The intertrial interval for each trial was 15 seconds. The day after the monkey completed the pre-training task, monkeys began a simple three-choice discrimination task. This task was administered to determine if there was a difference in the performance across the age groups in discriminating among three fixed stimuli based on reward contingency. The task presented the monkey with a pink square, orange cross and a brown 12-point star. The stimuli remained constant in terms of color and shape for each trial but appeared in a pseudo-random order in 9 different spatial locations on the screen for 80 trials per day. The pink square was the positive stimulus for all trials and all monkeys. A non-correctional procedure was used, such that the monkey was given a food reward only when he/she correctly touched the pink square on the screen. To reach criterion, the monkey had to choose the pink square on ten consecutive trials during one testing session.

Next, formal testing began on the CSST. Each day of testing consisted of 80 trials where three stimuli appeared in three of nine pseudo-random locations on the computer touch screen, as shown in **Figure 2**. The stimuli differed in two relevant dimensions, color (red, green, or blue) and shape (triangle, star, and circle). Each color and each shape was presented on every trial with all nine possible combinations of stimuli (i.e. red triangle, red star, red circle, blue triangle, etc.) presented in a pseudo-random sequence in a balanced fashion over four days of testing. If a monkey did not respond within one minute, the screen reverted to black, a non-response was recorded, and the intertrial interval began followed by the next trial.

**Figure 2.**
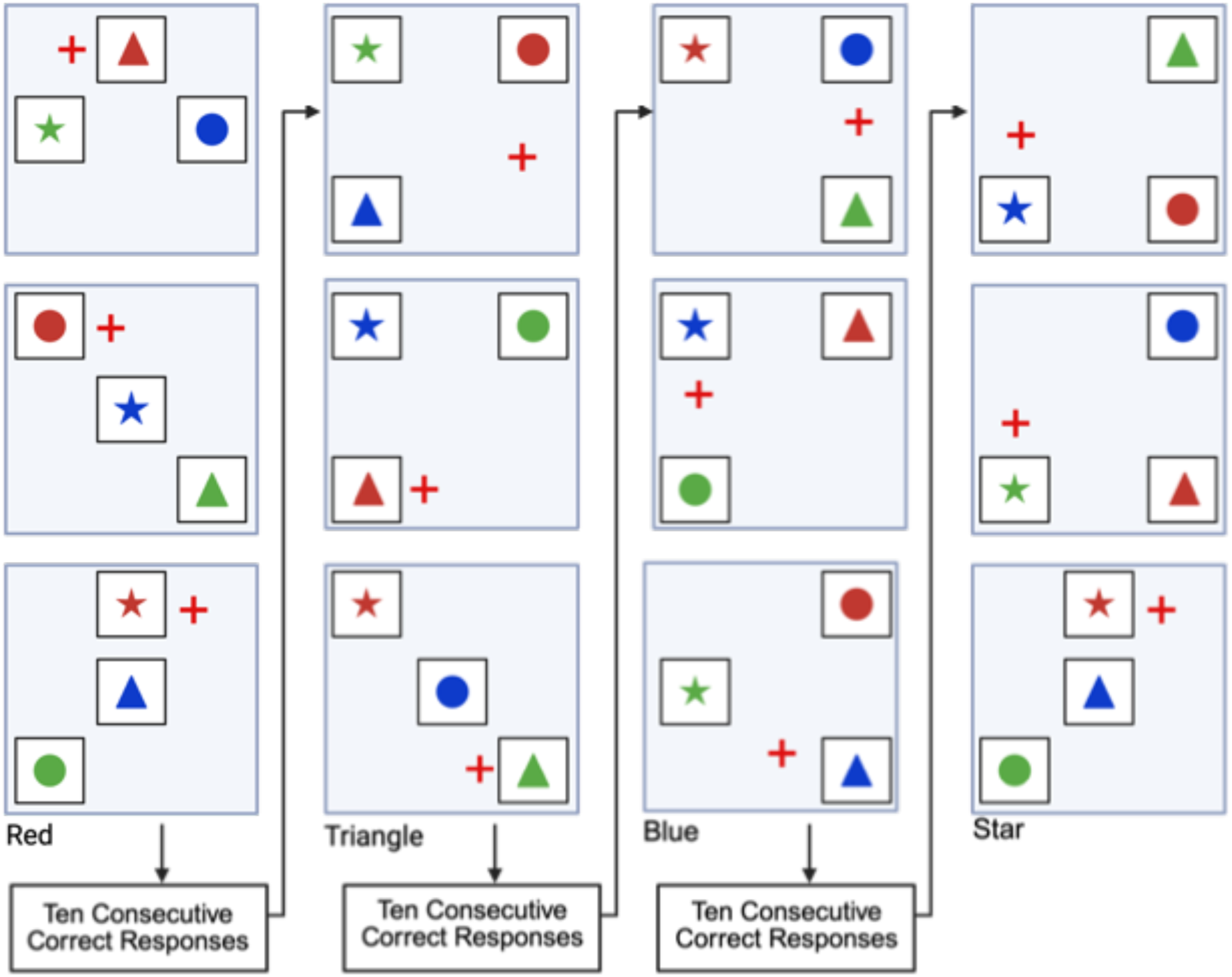
In this schematic of the Category Set Shifting Task (CSST) each blue screen (panel) represents one trial. On each trial the monkey is presented with three stimuli that vary in shape and color. During the first concept condition, the monkey must choose the red stimulus regardless of its shape as illustrated sequentially in the top three screens of this figure. Once the monkey chooses the correct stimulus on 10 consecutive trials, the computer switches the rewarded stimulus on the same testing day, without alerting the monkey. In the second concept condition, the monkey must choose the triangle shaped stimulus, regardless of the color as illustrated in the bottom three screens of the figure. Again, when the monkey chooses the correct stimulus for 10 consecutive trials the computer switches the rewarded stimulus on the same testing day, without alerting the monkey. Testing is continued in this same manner for the blue and star concept conditions.

For the first abstraction, red was designated as the positive dimension, and the monkey had to choose the red stimulus regardless of its shape to obtain a food reward. Once the monkey chose this stimulus on ten consecutive trials the program switched the rewarded contingency during the same testing session, without alerting the monkey. Now, the monkey had to choose the stimulus shaped like a triangle, regardless of its color, to obtain a food reward. Again, when the monkey reached a criterion of 10 consecutive responses, the computer switched the rewarded contingency so that the blue stimulus then had to be chosen, regardless of its shape, to obtain a food reward. Finally, when criterion was reached on the blue category, the contingency was switched to the last category, star.

A perseverative error was recorded when a monkey made an error by choosing a stimulus that contained a component of the previously rewarded concept. A broken set was recorded when a monkey achieved a span of six to nine consecutive correct responses and then made an error, missing criterion. A non-response was recorded when a monkey failed to touch the screen within one minute of stimulus presentation. Non-responses were also recorded as errors and reset the number of consecutive correct responses to zero.

### Outcome Measures and Data Analysis

All statistical analyses were performed in Matlab R2025b. To characterize relationships between cognitive performance and age across the adult lifespan, age was first analyzed as a continuous variable (i.e., age in years). Conventional categorical comparisons among young, middle-aged, and aged monkeys were then used as a complementary analysis to identify where broad age- group differences were detectable. To assess sex differences in age-dependent task performance, we included sex as well as the interaction of sex and age as regressors.

Robust linear regression (function ‘fitlm’ with ‘RobustOpts = On’) was used to analyze the relationship between behavioral performance and age as a continuous variable. This method uses iteratively reweighted least squares to downweight the impact of potential outlier data points. Then, to further analyze the effect of age, treated as a categorical variable (young, middle-aged and aged groups), on performance on the DNMS, DRST and CSST, we performed separate one-way analyses of variance (ANOVAs) (“anova” function in Matlab) for the outcome measures described above. In addition, a two-way repeated measures ANOVA, including age group and task condition, was performed for the trials, errors and perseverative errors on the three shift conditions (“fitrm” and “ranova” functions). After the ANOVAs, post hoc analyses were conducted using Tukey’s honestly significant difference procedure of estimated marginal means for multiple comparisons (“multcompare” function).

## Results

Continuous sampling across the adult lifespan allowed us to examine the extent to which age predicts changes in cognitive performance and whether different cognitive capacities show similar age-related trajectories. We first analyzed age as a continuous variable and then used conventional age-group comparisons as a complementary approach to identify where differences were detectable among young (5-12 years), middle- aged (13-20 years), and aged (>20 years) monkeys.

### Delayed Non-Matching to Sample

#### DNMS-Acq

Total trials and errors to criterion on the acquisition of the basic task were determined. The DNMS acquisition phase assessed age-related differences in rule learning, specifically development of the non- matching principle. With age as a continuous variable, we found significant increases in both the total number of trials (**Fig. 3A**; Table 3, F(1,88) = 8.98, p = 0.0035) and the number of errors (**Fig. 3B**; Table 3, F(1,88) = 16.1, p = 0.00012) required to reach criterion with increasing age. This is consistent with rule acquisition becoming progressively less efficient with increasing age. We also tested a regression model that included age, sex and their interaction as predictors and did not find statistical significance for sex (trials: p = 0.238, errors: p = 0.268) or the interaction between sex and age (trials: p = 0.098, errors: p = 0.120), indicating that the effect of age on learning performance did not differ significantly between males and females.

**Figure 3.**
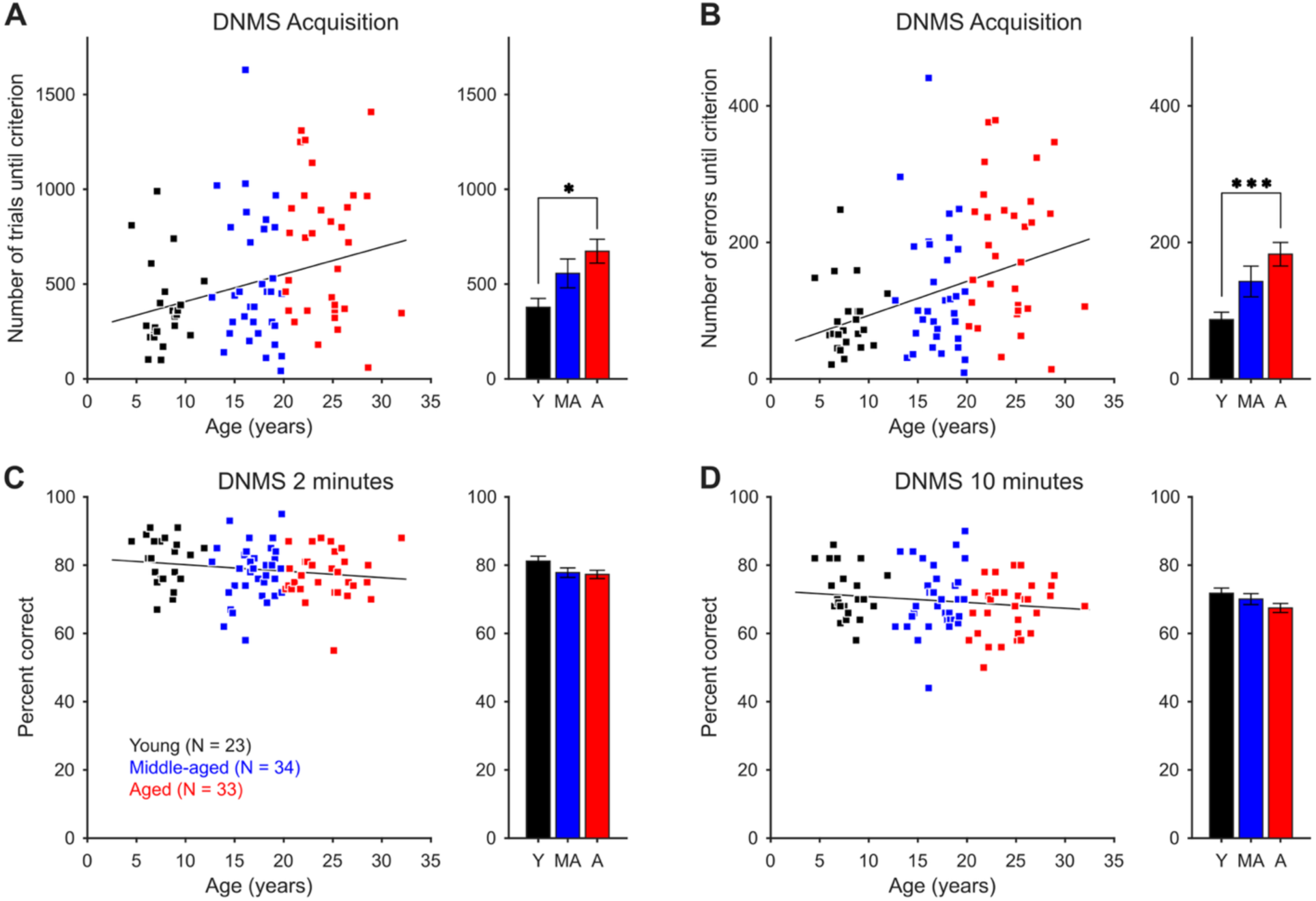
DNMS task acquisition and performance for longer delays across age. **A)** Left: Linear relationship of the total number of trials until criteria shows a significant correlation with age (*p* = 0.0068, n = 90 monkeys). Right: Average number of trials for the three age groups. Error bars indicate the SEM. **B)** Total number of errors until criteria vs. age (left panel; p = 0.00034, n = 90 monkeys) and mean ± SEM for the different age groups (right). **C)** DNMS performance for 2-minute delays vs. age. **D)** DNMS performance for 10-minute delays. See Table 3 for linear regression results and Table 4 for effect sizes and post hoc age group comparisons (**\*** = p ≤ 0.05; **\*\*** = p ≤ 0.01; **\*\*\*** = p ≤ 0.005).

**Table 3.** Robust linear regression analyses of age and measures of cognitive performance in the DNMS and DRST tasks.

| **Measurement** | **Equation** | ***R***<sup>**2**</sup> | **Adjusted *R*<sup>2</sup>** | ***F* statistic** | ***p* value** |
|---|---|---|---|---|---|
| DNMS acquisition total trials | *trials* = 14.4 • *age* + 264.2 | 0.0926 | 0.0823 | F(1.88) = 8.98 | 0.00354 |
| DNMS acquisition total errors | *errors* = 4.97 • *age* + 43.34 | 0.155 | 0.145 | F(1.88) = 16.1 | 0.000124 |
| DNMS 2 min – percent correct | *accuracy* = –0.189 • *age* + 82.0 | 0.031 | 0.0198 | F(1.88) = 2.8 | 0.0977 |
| DNMS 10 min – percent correct | *accuracy* = –0.171 • *age* + 72.5 | 0.0187 | 0.0075 | F(1.88) = 1.68 | 0.199 |
| DRSTsp span | *span* = –0.02 • *age* + 2.78 | 0.155 | 0.145 | F(1.88) = 16.2 | 0.00012 |
| DRSTob span | *span* = –0.051 • *age* + 3.98 | 0.214 | 0.205 | F(1.88) = 23.9 | 4.51 × 10<sup>-6</sup> |
| DRSTsp intercept of learning curve | *intercept* = –0.0027 • *age* + 2.17 | 0.0067 | –0.005 | F(1.88) = 0.593 | 0.443 |
| DRSTsp slope of learning curve | *slope* = –0.035 • *age* +1.16 | 0.149 | 0.14 | F(1.88) = 15.5 | 0.00017 |
| DRSTob intercept of learning curve | *intercept* = –0.027 • *age* +2.969 | 0.104 | 0.094 | F(1.86) = 10 | 0.00213 |
| DRSTob slope of learning curve | *slope* = –0.0370 • *age* +1.753 | 0.121 | 0.111 | F(1.86) = 11.8 | 0.000895 |

**Table 4.** Post hoc comparisons of age group differences across DNMS and DRST performance measures. Age groups: Y – young. MA – middle-aged. A – aged. The *p*-values were corrected for multiple comparisons.

| **Age Groups** | **Mean difference** | **95% confidence interval** | ***p*-value** | **Cohen’s *d*** |
|---|---|---|---|---|
| **DNMS acquisition total trials** |  |  |  |  |
| Y vs. MA | –178.7 | [–415.9. 58.5] | 0.1767 | –0.4739 |
| Y vs. A | –295.7 | [–534.3. –57.1] | 0.01111 | –0.9304 |
| MA vs. A | –117.0 | [–331.7. 97.7] | 0.3994 | –0.2849 |
| **DNMS acquisition total errors** |  |  |  |  |
| Y vs. MA | –55.9 | [–123.0. 11.1] | 0.1205 | –0.5170 |
| Y vs. A | –95.9 | [–163.4. –28.5] | 0.0030 | –1.133 |
| MA vs. A | –39.9 | [–100.6. 20.7] | 0.2644 | –0.3385 |
| **DRSTsp span** |  |  |  |  |
| Y vs. MA | 0.166 | [–0.039. 0.37] | 0.1351 | 0.4859 |
| Y vs. A | 0.355 | [0.155. 0.555] | 0.000164 | 1.1081 |
| MA vs. A | 0.189 | [0.001. 0.368] | 0.0363 | 0.6664 |
| **DRSTob span** |  |  |  |  |
| Y vs. MA | 0.4102 | [–0.125. 0.945] | 0.1664 | 0.4980 |
| Y vs. A | 0.7985 | [0.276. 1.321] | 0.00132 | 0.8894 |
| MA vs. A | 0.3883 | [–0.081. 0.858] | 0.12497 | 0.520 |

We next grouped the monkeys into categorical age groups (young, middle-aged, and aged) to complement the continuous-age analysis and identify whether differences were detectable among conventional age labels. A one-way ANOVA revealed a significant difference in age group performance (**Fig. 3A,B**; total trials: F(2,87) = 4.37, p = 0.0156; total errors: F(2,87) = 5.75, p = 0.0045). Post hoc tests (Table 4) showed a significant difference in total trials and errors to reach criterion between young and aged monkeys (Tukey’s post hoc test: young vs. aged groups, p = 0.011 for total trials and p = 0.0030 for total errors). However, there was no significant difference between young and middle-aged monkeys or between middle-aged and aged monkeys.

#### DNMS-Delays

Once the monkeys had reached criterion on the acquisition task, recognition memory performance was tested at two-minute and ten-minute delays. Accuracy did not significantly decline with age at either delay (**Fig. 3C**, two-minute delay, F(1,88) = 2.8, p = 0.10; and **Fig. 3D**, ten-minute delay, F(1,88) = 1.68, p = 0.20). Moreover, a one-way ANOVA comparing age groups also did not show significant differences (**Fig. 3C**, two-minute delay, F(2,87) = 2.14, p = 0.12; and **Fig. 3D**, ten-minute delay, F(2,87) = 1.97, p = 0.145). These results are in contrast with previous reports of a significant age dependence of short-term memory performance on the DNMS delay task. ^4, 5^ These differences may be due, in part, to different testing paradigms in each study. In our studies, monkeys completed the DNMS task first in the testing battery and after making criterion on the basic task (10sec delay), they then completed 10 trials/day for 10 ten days (100 trials in total) for each of 2- and 10-minute delays. In Baxter et al, 2023, animals completed the Delayed Response Task prior to testing on the DNMS task which provided the animals with more experience and therefore may account for the difference in performance across tasks. Further, for DNMS delays, the animals in Baxter et al, 2023 completed 100 trials, 20/day at 2-minute delays and 50 trials, 5/day at 10 minute delays.

### Delayed Recognition Span Task – Spatial and Object

#### DRST

For both the DRST spatial and object tasks, the mean total span achieved by each monkey was determined across the 10 trials per day for 10 days of testing (total of 100 trials for each task) (**Fig. 4A,B**). To examine practice-related changes in performance across sessions, the mean span was calculated for each block of ten trials, corresponding to each daily testing session. Finally, a linear regression was fit across sessions for each monkey to quantify initial performance (intercept) and the rate of learning across sessions (slope) (**Fig. 5C-F**). In supplementary analysis the mean of all monkeys in each age group for each trial was also calculated.

**Figure 4.**
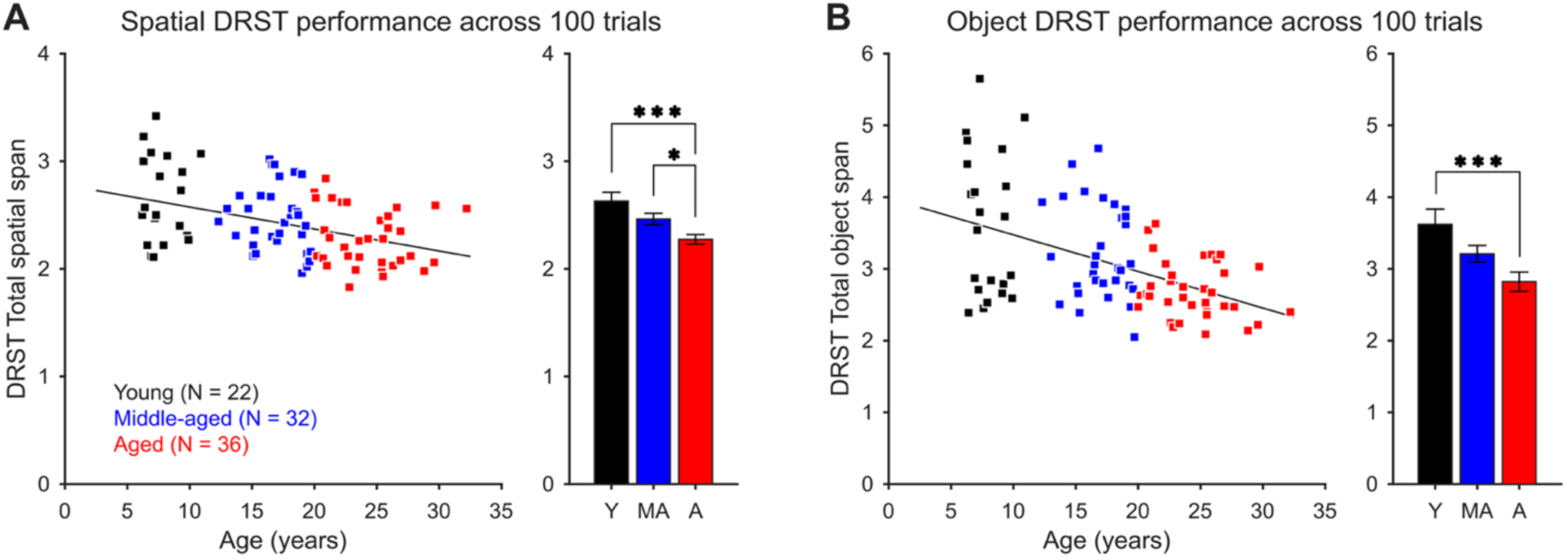
Overall task performance in the spatial and object DRST across age. **A**) Left: Relationship between age and trial-averaged DRST spatial span (*p* = 0.00013). Right: Mean ± SEM DRST spatial span in young, middle-aged and aged monkeys. **B)** Left: Trial-averaged performance in the object DRST task (p = 6.85 × 10^-6^). Right: performance for different age-groups. See Table 3 for linear regression results and Table 4 for effect sizes and post hoc age group comparisons (**\*** = p ≤ 0.05; **\*\*** = p ≤ 0.01; **\*\*\*** = p ≤ 0.005).

**Figure 5.**
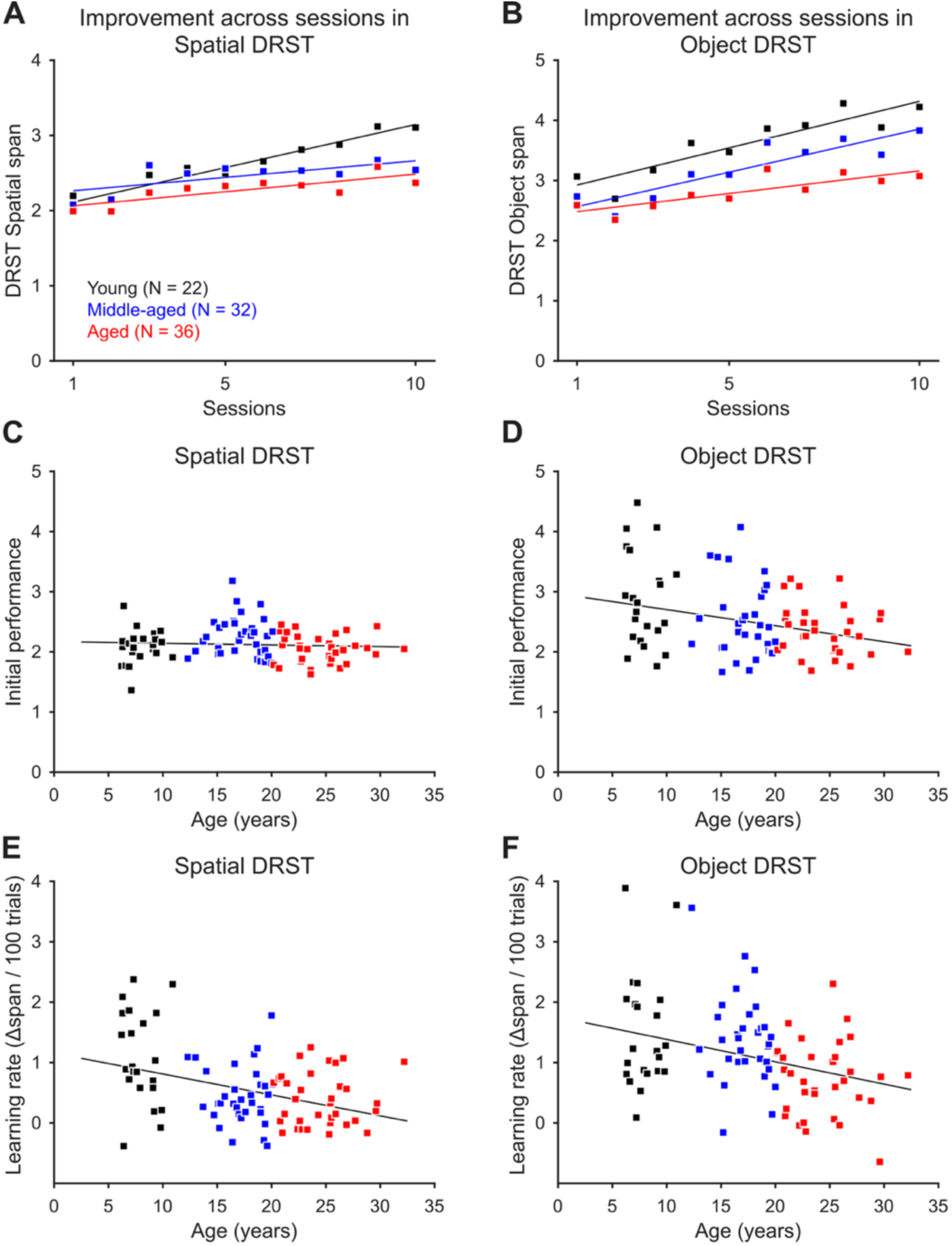
Learning across sessions in the spatial DRST and the object DRST. **A, B)** Mean DRST span across all monkeys of the corresponding age group for each of 10 consecutive testing sessions. **C, D**) Initial performance (intercept of learning curve) vs. age for the spatial DRST (**C**; *p* = 0.53) and the object DRST (**D**; *p* = 0.0051). **E, F)** Learning rate (slope of the learning curve) vs. age for spatial DRST (*p* = 0.00020) and object DRST (*p* = 0.0013). See Table 3 for linear regression results and Table 5 for mixed-effects model results.

**Table 5.** Fixed effects from a linear mixed-effects model predicting DRST performance (block span) across testing blocks. for the spatial and object versions of the task. Age groups: Y – young. MA – middle-aged. A – aged.

| **Predictor** | **Estimate** | **SE** | **t** | **df** | **p-value** | **95% CI (Lower–Upper)** |
|---|---|---|---|---|---|---|
| **DRST-Spatial Task** |  |  |  |  |  |  |
| Y | 1.998 | 0.063 | 31.71 | 894 | < 10<sup>−16</sup> | [1.875. 2.122] |
| MA vs. Y | 0.208 | 0.084 | 2.50 | 894 | 0.0127 | [0.045. 0.372] |
| A vs. Y | 0.019 | 0.076 | 0.25 | 894 | 0.802 | [–0.130. 0.169] |
| Session | 0.115 | 0.016 | 7.05 | 894 | 3.65 × 10<sup>−12</sup> | [0.083. 0.147] |
| MA × Session interaction | –0.066 | 0.018 | –3.63 | 894 | 0.0003 | [–0.101. –0.030] |
| A × Session interaction | –0.072 | 0.018 | –3.99 | 894 | 7.03 × 10<sup>−5</sup> | [–0.108. –0.037] |
| **DRST-Object Task** |  |  |  |  |  |  |
| Y | 2.731 | 0.160 | 17.06 | 894 | < 1 × 10<sup>−16</sup> | [2.417. 3.046] |
| MA vs. Y | –0.198 | 0.226 | –0.87 | 894 | 0.382 | [–0.642. 0.246] |
| A vs. Y | –0.396 | 0.178 | –2.22 | 894 | 0.0265 | [–0.745. –0.046] |
| Session | 0.151 | 0.021 | 7.17 | 894 | 1.58 × 10<sup>−12</sup> | [0.110. 0.193] |
| MA × Session interaction | –0.016 | 0.024 | –0.65 | 894 | 0.514 | [–0.063. 0.032] |
| A × Session interaction | –0.067 | 0.024 | –2.75 | 894 | 0.0062 | [–0.115. –0.019] |

For the DRST task, the mean span across ten days of testing for both the spatial and object versions of the task was determined for each monkey. We found a gradual age-related decline in working memory performance in both the spatial and the object DRST tasks (**Fig. 4A,B**). Robust linear regression of mean DRSTsp and DRSTob total span showed that age has a significant effect on working memory (Table 3; DRSTsp: F(1,88) = 16.2, p = 0.00012; DRSTob, F(1,88) = 23.9, p = 4.51 × 10^-6^). Adding sex and the interaction of age and sex to the regression model did not identify significant effects of sex (DRSTsp: p = 0.646, DRSTob: p = 0.400) or the interaction between sex and age (DRSTsp: p = 0.526, DRSTob: p = 0.236). A one-way ANOVA age group analysis revealed a significant difference between age groups (DRSTsp: F (2,87) = 9.3, p = 0.0002; DRSTob: F(2,87) = 6.75, p = 0.0019). Tukey’s post-hoc test results (Table 4) showed a significant difference between young and aged monkeys on both DRSTsp and DRSTob (DRSTsp: p = 0.00016; DRSTob: p = 0.0013) and between middle-aged and aged monkeys on the DRSTsp (DRSTsp: p = 0.036; but not DRSTob, p = 0.12).

In sum, age-related changes in task performance were similar in both the spatial and object versions of this task, despite overall higher scores on the object version. Moreover, the mean spans obtained on each version of the task were highly correlated with each other even when factoring out the common age-dependence (partial correlation taking into account age: r = 0.48, p = 1.84 × 10^-6^).

We also examined how performance changed across testing of repeated DRST sequences to assess how age affected practice mediated improvement. We characterized this learning effect by averaging the DRST span within blocks of 10 trials across 10 days of testing for each age group. This analysis revealed an age-dependent increase in performance over time, with the greatest improvement in young monkeys for both tasks (**Fig. 5A,B**). This can also be observed when plotting the average span for each monkey against age in the different testing sessions (**Suppl. Fig. 1** and **Suppl. Fig. 2**). To quantitatively assess how repeated testing influenced recognition span performance in both the spatial and object versions of the DRST, we fit linear mixed-effects models for each task. Each model included age group (young, middle-aged, aged), block number, and their interaction as fixed effects, with random intercepts and slopes by subject. The results are summarized in Table 5. For the DRSTsp task (**Fig. 5A**), block span performance of young monkeys improved significantly across sessions (p = 3.65 × 10^−12^), consistent with a learning effect. Both middle-aged (p = 0.0003) and aged animals (p = 7.03 × 10^−5^) exhibited significantly less improvement across sessions compared to young monkeys suggesting less efficient learning across trials. For the DRSTob task (**Fig. 5B**), young monkeys again improved over sessions (p = 1.58 × 10^−12^). Middle-aged animals did not differ significantly from young animals in their learning rate (p = 0.514). However, aged monkeys again showed less efficient learning across trials (p = 0.0062).

We further analyzed the learning effect across age by calculating a learning curve for each individual monkey, characterized by initial performance in the first block of trials (intercept) and the rate of improvement across blocks (slope) (**Fig. 5C-F**). Robust linear regressions for DRSTsp, with age as a continuous variable, showed no systematic age-dependence of the initial performance (**Fig. 5C**; F(1,88) = 0.593, p = 0.443). However, in the DRSTob task, the initial performance decreased with age (**Fig. 5D**; F(1,86) = 10, p = 0.002). Further, the slope of the learning curves in both tasks significantly decreased with age (DRSTsp: **Fig. 5E**; F(1,88) = 15.5, p = 0.0002; DRSTob: **Fig. 5F**; F(1,86) = 11.8, p = 0.0009) confirming that aged monkeys improved less efficiently across repeated testing.

Together, these results indicate that age-related decline reflects not only reduced working-memory capacity, but also a diminished ability to improve performance through repeated experience. They also suggest earlier vulnerability of practice-related improvement in the spatial task, where attenuation was already detectable in middle-aged monkeys.

#### Category Set Shifting Task

The total number of trials and errors to criterion for the three-choice discrimination task, the total number of trials and errors to criterion for the red condition (initial abstraction) and the total number of trials, errors and perseverative errors for the three shift conditions (triangle, blue and star) were determined. Perseverative errors were also calculated as a percentage of total shift trials to account for differences in the number of opportunities to perseverate.

We first examined CSST performance using linear regression models with age as a continuous variable. This revealed a slight but significant increase in the number of trials and errors required to reach criterion with advancing age during initial abstraction of the category rule (**Suppl. Fig. 3**; Table 6; trials: F(1,89) = 6.23, p = 0.0144; errors: F(1,89) = 4.59, p = 0.0349). We also replicated the previously reported substantial age-dependent increases in total trials and errors across the three shift conditions, as well as in total perseverative errors (Table 6^10^). To assess potential sex differences, we extended the regression models to include sex and the age × sex interaction as additional regressors. No significant sex effects were observed for the number of trials (sex: p = 0.938; age × sex interaction: p = 0.653) or errors (sex: p = 0.820; age × sex interaction: p = 0.549) across the three shift conditions, nor for total perseverative errors (sex: p = 0.890; age × sex interaction: p = 0.663). Collectively, these findings point to an age-related decline in both abstraction of the initial category rule and cognitive flexibility following category switching, with no evidence of sex differences.

**Table 6.** Robust linear regression analyses of age and measures of cognitive performance in the CSST tasks.

| **Measurement** | **Equation** | ***R*<sup>2</sup>** | **Adjusted *R*<sup>2</sup>** | ***F* statistic** | ***p* value** |
|---|---|---|---|---|---|
| CSST Total trials for first category (red) | *trials* = 4.72 • *age* + 190.4 | 0.0654 | 0.0549 | F(1.89) = 6.23 | 0.0144 |
| CSST Total errors for first category (red) | *errors* = 1.77 • *age* + 85.78 | 0.049 | 0.0383 | F(1.89) = 4.59 | 0.0349 |
| CSST Total trials for three shift conditions | *trials* = 32.11 • *age* + 682.8 | 0.255 | 0.247 | F(1.89) = 30.5 | 3.3 × 10<sup>-7</sup> |
| CSST Total errors for three shift conditions | *errors* = 15.9 • *age* + 380.4 | 0.262 | 0.253 | F(1.89) = 31.6 | 2.19 × 10<sup>-7</sup> |
| CSST Total perseverative errors | *p-errors* = 16.4 • *age* + 242.4 | 0.244 | 0.236 | F(1.89) = 28.8 | 6.31 × 10<sup>-7</sup> |
| CSST Total perseverative errors as a % of shift trials | *p-errors* = 0.0698 • *age* + 40.3 | 0.0074 | -0.0037 | F(1.89) = 0.669 | 0.416 |
| CSST Total number of broken sets | *broken sets* = 0.326 • *age* + 6.36 | 0.132 | 0.122 | F(1.89) = 13.6 | 0.00040 |
| CSST Total non-responses | *non-responses* = 1.96 • *age* + 30.44 | 0.234 | 0.225 | F(1.89) = 27.1 | 1.21 × 10<sup>-6</sup> |

To complement the continuous-age analyses, we also compared performance across the three conventional age groups. Middle-aged and aged monkeys required more trials on average to reach criterion during the initial abstraction phase (**Fig. 6A**), although the group means did not differ significantly due to substantial inter-subject variability (ANOVA: F(2,88) = 2.69, p = 0.073). Likewise, the total errors to criterion for the initial abstraction did not significantly differ across groups (F(2,88) = 1.526, p = 0.223).

**Figure 6.**
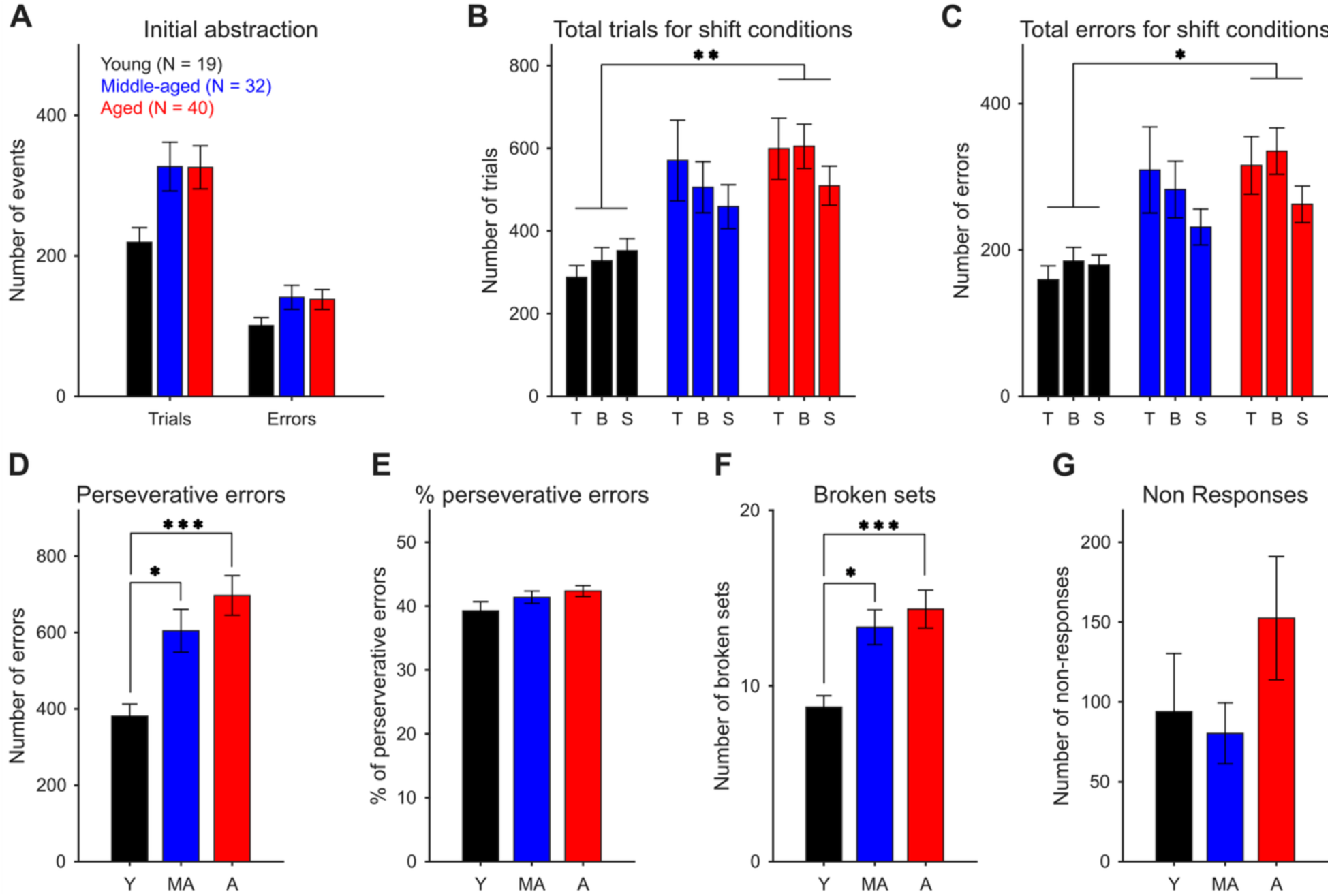
Comparison of CSST performance between age groups (young, middle-aged and aged monkeys). **A)** Number of trials and errors in the initial abstraction task (red condition). **B)** Total trials in each of the three shift conditions. **C)** Total errors in each of the three shift conditions. **D)** Total perseverative errors across the three shift conditions. **E)** Perseverative errors as percent of shift trials. **F)** Total broken sets. **G)** Number of non- responses. Errorbars indicate the mean ± SEM. See Table 7 for effect sizes and post hoc age group comparisons (**\*** = p ≤ 0.05; **\*\*** = p ≤ 0.01; **\*\*\*** = p ≤ 0.005)

**Table 7.**
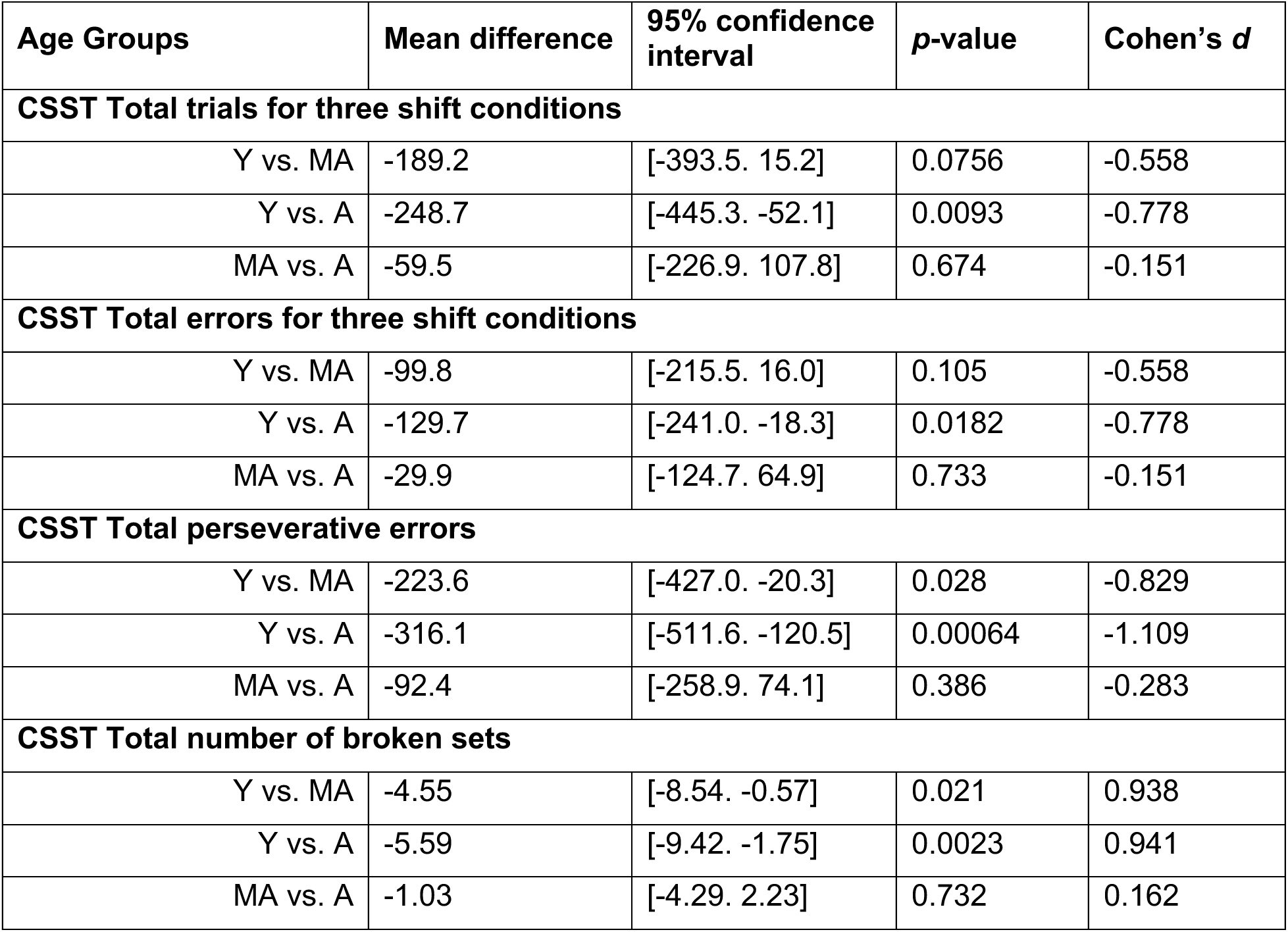
Post hoc comparisons of age group differences across CSST performance measures. Age groups: Y – young. MA – middle-aged. A – aged. The *p*-values were corrected for multiple comparisons.

| **Age Groups** | **Mean difference** | **95% confidence interval** | ***p*-value** | **Cohen’s *d*** |
|---|---|---|---|---|
| **CSST Total trials for three shift conditions** |  |  |  |  |
| Y vs. MA | -189.2 | [-393.5. 15.2] | 0.0756 | -0.558 |
| Y vs. A | -248.7 | [-445.3. -52.1] | 0.0093 | -0.778 |
| MA vs. A | -59.5 | [-226.9. 107.8] | 0.674 | -0.151 |
| **CSST Total errors for three shift conditions** |  |  |  |  |
| Y vs. MA | -99.8 | [-215.5. 16.0] | 0.105 | -0.558 |
| Y vs. A | -129.7 | [-241.0. -18.3] | 0.0182 | -0.778 |
| MA vs. A | -29.9 | [-124.7. 64.9] | 0.733 | -0.151 |
| **CSST Total perseverative errors** |  |  |  |  |
| Y vs. MA | -223.6 | [-427.0. -20.3] | 0.028 | -0.829 |
| Y vs. A | -316.1 | [-511.6. -120.5] | 0.00064 | -1.109 |
| MA vs. A | -92.4 | [-258.9. 74.1] | 0.386 | -0.283 |
| **CSST Total number of broken sets** |  |  |  |  |
| Y vs. MA | -4.55 | [-8.54. -0.57] | 0.021 | 0.938 |
| Y vs. A | -5.59 | [-9.42. -1.75] | 0.0023 | 0.941 |
| MA vs. A | -1.03 | [-4.29. 2.23] | 0.732 | 0.162 |

A two-way ANOVA on total trials to criterion across the three shift conditions, with age group as a between- subjects factor and condition as a repeated measure, revealed a significant main effect of age group (F(2,88) = 4.60, p = 0.0126). Neither the main effect of condition (p = 0.42) nor the age group × condition interaction (p = 0.38) reached significance. Tukey’s post hoc comparisons identified a significant difference between young and aged monkeys (p = 0.0093), while middle-aged monkeys did not differ significantly from either young (p = 0.0756) or aged (p = 0.674) animals (**Fig. 6B**; Table 7). A parallel analysis of total errors to criterion across the three shift conditions yielded a significant main effect of age group (F(2,88) = 3.90, p = 0.024), with no significant effect of condition (p = 0.098) or age group × condition interaction (p = 0.42). Post hoc testing again revealed a significant difference between young and aged monkeys (p = 0.018), whereas middle-aged monkeys did not differ significantly from young (p = 0.105) or aged (p = 0.733) groups (**Fig. 6C**; Table 7). Thus, both trials and errors to criterion across shift conditions consistently distinguished aged from young monkeys, with middle-aged animals showing intermediate, performance that was in the same direction but non-significant.

For total perseverative errors across all shift conditions, a one-way ANOVA revealed a significant main effect of age group (**Fig. 6D**; F(2,88) = 7.43, p = 0.0010). Tukey’s post hoc tests showed that both middle-aged (p = 0.028) and aged (p = 0.00064) monkeys committed significantly more perseverative errors than young animals. However, because middle-aged and aged monkeys required more trials to reach criterion, they had greater opportunity to perseverate. When perseverative errors were therefore expressed as a percentage of shift trials, no significant differences across age groups were detected (**Fig. 6E**; F(2,88) = 1.99, p = 0.142). The increase in raw perseverative errors is therefore best interpreted as part of a broader decline in task efficiency rather than as evidence for a disproportionate tendency to perseverate. Analysis of total broken sets revealed a significant age-group effect (**Fig. 6F**; F(2,88) = 6.22, p = 0.0030), with younger monkeys showing more consistent response patterns, reflected in fewer broken sets, compared to middle-aged and aged animals (**Fig. 6F**; Table 7). Finally, total non-responses did not differ significantly across age groups (**Fig. 6G**; F(2,88) = 1.45, p = 0.24).

Taken together, these results demonstrate progressive age-related declines in abstraction and set-shifting. Set- shifting significantly distinguished aged from young monkeys, while middle-aged monkeys showed intermediate performance, suggesting that decline begins prior to old age. The apparent increase in perseverative errors with age is largely attributable to the greater overall number of trials and errors in older animals, rather than a disproportionate tendency to perseverate. The overall pattern of CSST performance across age groups closely mirrors the age-related deficits observed in middle-aged and elderly humans on the Wisconsin Card Sorting Task ^21^, the paradigm the CSST was designed to model in non-human primates.

Across the behavioral battery, continuous-age analyses revealed graded changes in multiple aspects of cognitive performance, whereas categorical comparisons most often identified differences only between the endpoints of adulthood. Although the magnitude of these relationships varied across measures, age-related decline was evident in both core cognitive capacities and the ability to improve or adapt behavior with experience.

## Discussion

Cognitive aging in the rhesus monkeys in the present study is expressed as a gradual shift in population-level performance across the adult lifespan. The continuous relationships observed between chronological age and cognitive performance extend the established observation that young and aged animals differ substantially from one another. Across the behavioral battery in this study, age predicted both reduced cognitive capacity and a diminished ability to improve performance through experience. However, the strength and timing of these relationships varied across measures, indicating that cognitive aging is not expressed uniformly across domains.

This pattern of cognitive decline was most clearly resolved by the continuous-age analyses presented here. Categorical comparisons provided a coarser but complementary view of these relationships. Age-group differences most often distinguished young from aged animals, consistent with the accumulation of gradual declines across adulthood. The categorical differences are therefore most parsimoniously interpreted as lower- resolution manifestations of broader lifespan gradients, rather than as evidence for sharply bounded stages of decline. Specifically, increasing age was associated with poorer rule acquisition, working-memory capacity, practice-related improvement, abstraction, and set-shifting, while recognition-memory performance at extended delays was comparatively preserved. For selected measures, the accumulated change was sufficiently pronounced to distinguish middle-aged animals from one endpoint: practice-related improvement on the spatial DRST was already attenuated in middle age, whereas spatial working-memory capacity declined further between middle and advanced age. Measures of response consistency on the CSST also distinguished middle-aged from young animals. No significant sex differences in age-related cognitive performance were detected in this cohort.

This graded decline in performance with increasing age was revealed by the inclusion of a substantial number of middle-aged animals. Prior comparisons of young and aged monkeys accurately identified the large cognitive differences associated with advanced age.^5, 6, 9, 22^ Rather than forming a sharply distinct behavioral class, middle- aged monkeys frequently exhibited intermediate levels of performance. Their inclusion therefore does more than fill a missing category between two well-defined endpoints, it reveals a population-level trajectory of progressive and graded cognitive declines across adulthood. Endpoint comparisons therefore capture the magnitude of cognitive impairment in aged animals, but they incompletely describe the broader process of cognitive aging that is revealed with the inclusion of middle-aged monkeys.

The prevalence of intermediate performance values across middle age indicates that cognitive aging in rhesus monkeys is not a matter of increasing the likelihood of stepwise transition from intact to severely impaired performance. Instead, middle-aged animals are more likely to have partially degraded performance, suggesting that progressive population-level change includes gradual cognitive decline within the context of the wide inter- individual variability. However, this inference must be distinguished from a stronger claim that cannot be established from our cross-sectional data. The present study does not show that every individual monkey declines gradually over time. Individual animals may follow different trajectories with some deteriorating slowly and gradually over time while others may undergo abrupt changes. Individual differences in neurobiological vulnerability may alter the timing, domain-specificity, and severity of decline. This hypothesis is supported by the findings of Vanderlip et al. (2025)^23^ who similarly observed distinct learning curves in rhesus monkeys across the life span on object DRST. However, to fully understand these trajectories, longitudinal studies are required to determine the shape of within-animal trajectories and to identify the factors that confer resilience or vulnerability.

### Age-Related Cognitive Decline in Humans

The pattern of cognitive aging observed in the current study demonstrates parallels between rhesus monkeys and humans, supporting the translational relevance of the rhesus monkey as a model of age-related cognitive decline. Normal human cognitive aging is increasingly recognized as a gradual, domain-specific process that begins during middle adulthood and progresses across the lifespan, with substantial inter-individual variability.^24–29^ Specifically, studies of healthy human aging have consistently demonstrated that cognitive function does not decline in a uniform fashion across domains and is not restricted to the oldest old. Changes occur in learning, memory, processing speed, and executive function at different stages of adulthood (Harada et al., 2013; Hartshorne and Germine, 2015; Juan and Adlard, 2019). Further, executive functions, cognitive flexibility, working memory, processing speed, and the acquisition of new information are the earliest and most consistently age-affected domains,^24, 27, 28^ whereas crystallized knowledge and aspects of long-term semantic memory remain relatively preserved until later in life.^25, 28^ Importantly, the earliest detectable changes often involve reduced efficiency in acquiring new information and adapting behavior, rather than complete loss of previously learned abilities.^27, 28, 30^

Hartshorne and Germine (2015) demonstrated that different cognitive abilities reach peak performance at distinct ages, with some declining in young adulthood while others remain stable or continue to improve into middle and later adulthood. Similarly, Harada et al. (2013) reported that working memory, novel learning, and executive function, are particularly vulnerable to aging, whereas crystallized abilities are relatively preserved. Recently in a large-scale study, Matsui et al. (2026), examining more than 300 cognitively normal adults aged 60 years and older, demonstrated that processing speed and executive function showed significant impairments by the early 70s, preceding more pronounced changes in other cognitive abilities. Importantly, these findings demonstrate that measurable cognitive impairments occur even in individuals clinically classified as cognitively normal, reinforcing the distinction between normal cognitive aging and dementia.

Consistent with these studies with aging humans, our analysis of cognitive function in rhesus monkeys spanning the adult lifespan demonstrated progressive, domain-dependent declines in rule acquisition, working-memory capacity, cognitive flexibility, and the ability to improve performance with experience. Further, the inclusion of a large middle-aged group and the use of continuous-age analyses revealed that age-related cognitive differences were evident before advanced age and are not uniformly expressed across cognitive domains. The decline in practice-dependent improvement observed in our monkeys is also consistent with human studies demonstrating reduced efficiency in acquiring new information with advancing age.

Taken together, our findings and the results of studies with aged humans, suggest that normal cognitive aging represents a gradual, multidimensional process with conserved features in both monkeys and humans, rather than a uniform deterioration progressing with age. Although the timing and magnitude of age-related changes differs across cognitive domains and species, the convergence of these findings underscores the value of rhesus monkeys for investigating the neurobiological mechanisms underlying early cognitive vulnerability.

### Neurobiological Basis of Age-Related Cognitive Decline

The degradation of cognitive performance across aging is consistent with the aggregate burden of neurobiological changes that unfold over individual timelines. Although widespread neuronal loss is not a defining feature of normal aging,^31–34^ circuits supporting cognition undergo remodeling across multiple levels of neural organization. Local circuits are altered at the level of individual neurons, as aging changes intrinsic cellular physiology in the hippocampus^35, 36^ and prefrontal cortex^37–40^ and results in loss and reorganization of synaptic architecture in the prefrontal cortex,^32, 38, 41–43^ hippocampus^44, 45^ and V1.^46^ These local vulnerabilities are compounded by degradation of the long-range myelinated pathways that mediate cognition across frontal, parietal, and temporal networks^47^, and this white-matter degeneration can be exacerbated by age-related changes in glial and inflammatory regulation.^48–52^ Also, altered neuromodulatory input with age modifies and dysregulates the function of cognitive circuits.^1, 53–59^ Several of these changes are associated with degraded behavioral performance,^38, 39, 43, 44, 55, 60^ while the functional consequences of others may become apparent only under more sensitive or domain-specific assessments. Each biological consequence of aging may provide contributions to distinct trajectories of cognitive vulnerability. Variation in the timing, severity, anatomical distribution, and degree of compensation across these processes may therefore underlie the individual differences documented here and help explain why cognitive capacities need not decline in lockstep. Future studies will be required to determine how distinct patterns of biological aging shape unique profiles of cognitive decline.

### Conclusions

The combination of gradual population-level change and marked inter-individual variability observed in this study closely parallels a central challenge in the study of human cognitive aging. In humans, cognitive trajectories are heterogeneous and difficult to disentangle from subclinical pathology, health history, lifestyle, and environmental exposures. Rhesus monkeys provide an opportunity to study these processes under more controlled conditions and to follow individuals longitudinally across adulthood. The present results establish an age-defined behavioral baseline against which individual deviations and their biological correlates can be interpreted. This framework provides a foundation for determining why some individuals remain resilient while others become vulnerable, and ultimately for evaluating whether interventions can preserve cognitive function before advanced impairment develops.

## Acknowledgments

The authors thank Bethany Bowley, Penny Shultz, Karen Slater, Rudolph Beiler, DVM and Ethan Gaston for their invaluable assistance with this project and the CERCA Program/Generalitat de Catalunya for institutional support. This project was funded by: NIH/NIA RF1AG043640, NIH/NIA RF1AG062831, NIH/NIA RF1 AG068168, NIH/NIA P01 AG000001, NIA-NSF CRCNS R01 AG071230, as well as by the Agencia Estatal de Investigación (AEI) of the Spanish Ministry of Science, Innovation and Universities through projects PCI2020- 112035 / AEI / 10.13039/501100011033, PID2023-147627OB-I00 / AEI / 10.13039/501100011033, and the Severo Ochoa and María de Maeztu Program for Centers and Units of Excellence in R&D (CEX2020-001084- M).

---

**Supplementary Figure 1.**
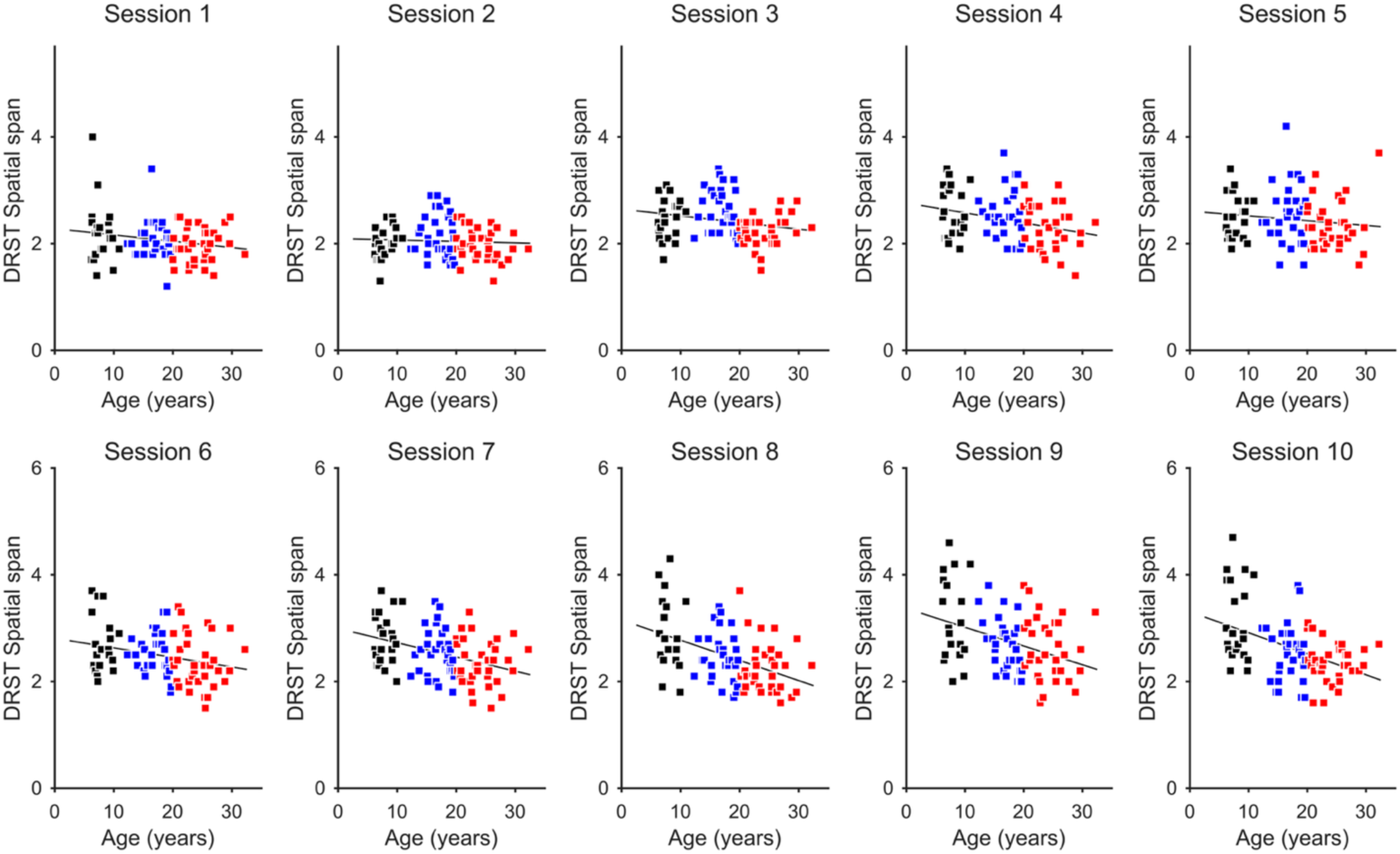
Average DRST spatial span vs. age in the 10 consecutive sessions for each monkey (n = 90).

**Supplementary Figure 2.**
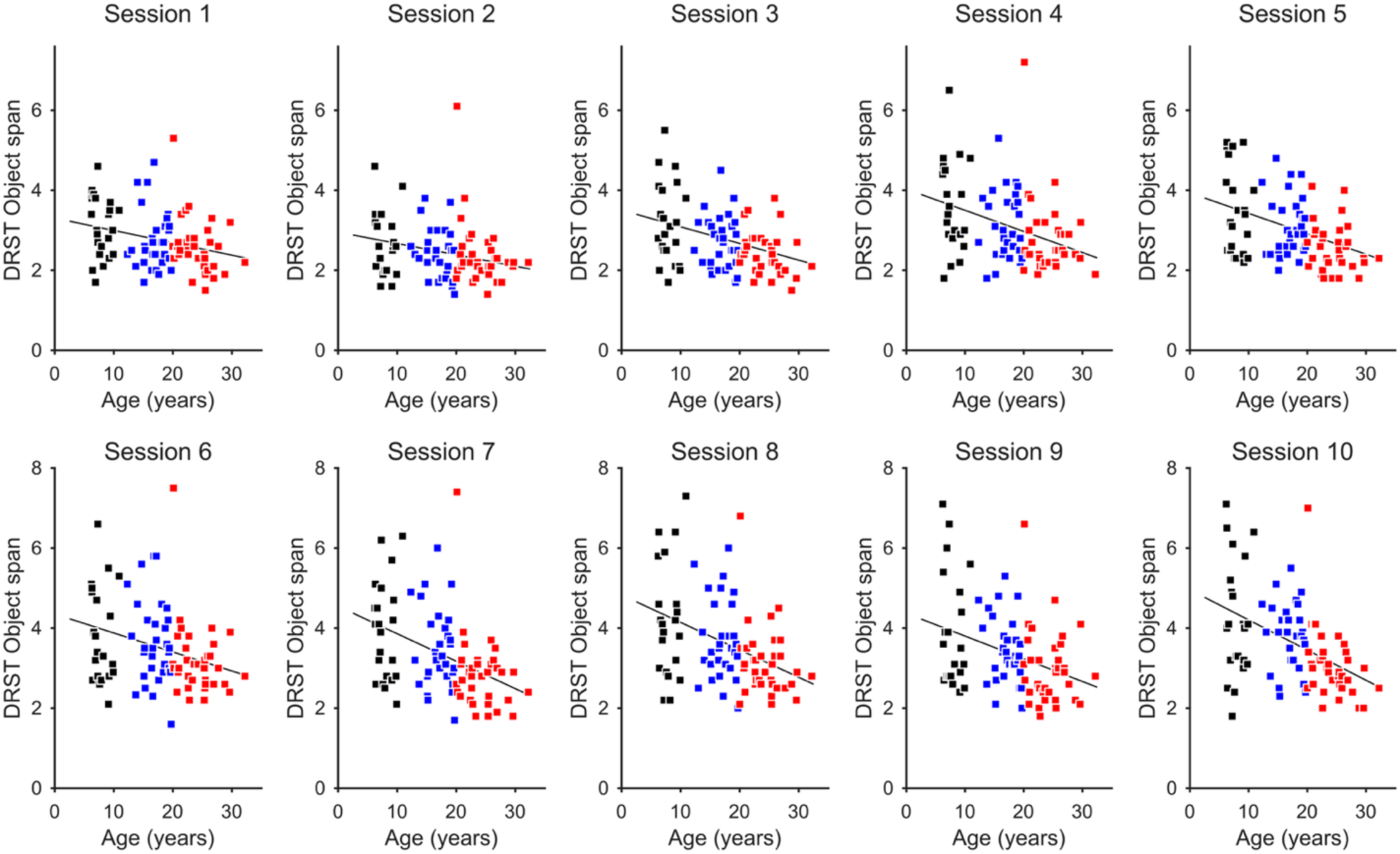
Average DRST object span vs. age in the 10 consecutive sessions for each monkey (n = 90).

**Supplementary Figure 3.**
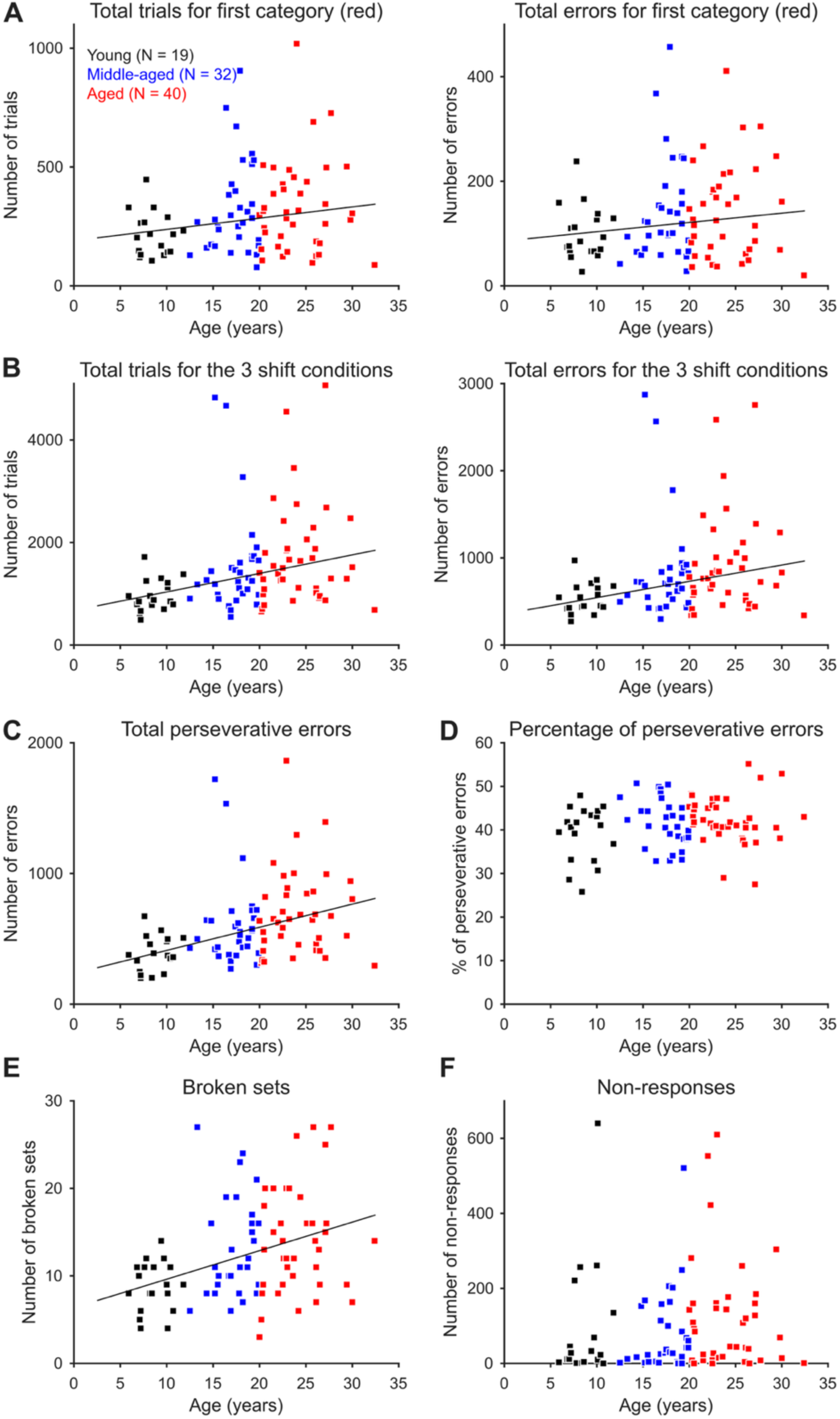
Relationship of CSST performance and age. Regression lines were computed with robust linear regression and are shown for the cases in which there is a significant relationship. See Table 6 for the statistical results. **A)** Total trials and errors in the initial abstraction (red condition) vs. age. **B)** Total trials and errors in the three shifts across condition vs. age. **C**) Total perseverative errors vs. age. **D)** Perseverative errors as a percentage of shift trials vs. age. **E)** Number of total broken sets across conditions vs. age. **F)** Number of non-responses vs. age.

